# Suppression of early pro-inflammatory senescent signature post-radiotherapy mitigates chronic bone damage

**DOI:** 10.64898/2026.03.01.708630

**Authors:** David Achudhan, Jacob Orme, Ritika Sharma, Aqsa Komel, Khubaib Gul Khan, Thomas A. White, Nathan K. LeBrasseur, Sundeep Khosla, Sean S. Park, Robert J. Pignolo, Abhishek Chandra

**Author notes:** **<u>Corresponding author</u>:** Abhishek Chandra, Ph.D., M.S., Guggenheim 7, Mayo Clinic College of Medicine, 200 First Street SW, Rochester, MN 55905.

## Abstract

Cellular senescence has been implicated in the pathophysiology of radiotherapy-related bone loss. Based on our previous work, clearance of senescent cells using genetic and pharmacological tools alleviates the anomalies associated with radiation-associated bone deterioration. The pro-inflammatory senescence associated secretome referred to as senescence associated secretory phenotype (SASP), is a hallmark of cellular senescence. The modulation of SASP by senomorphic drugs, potentially can suppress the pro-inflammatory secretome of senescent cells, irrespective of the underlying senescence mechanism. In this study we tested a senomorphic drug, ruxolitinib, a Janus kinase inhibitor (JAKi), during acute and chronic radiotherapy related effects on the bone. Our clinical data indicate an early increase in several pro-inflammatory SASP proteins following radiotherapy of spinal metastasis in prostate cancer patients. Longitudinal assessment of SASP-related genes confirmed this acute elevation in several SASP markers in systemic circulation following irradiation of mouse femurs. In a proof-of-concept study, following two preclinical radiotherapy regimens of cumulative doses of 30Gy (5 x 6Gy) and 60Gy (5 x 12Gy), a senomorphic approach of JAKi treatment was more effective in alleviating radiation-related bone loss compared to the senolytic cocktail of D+Q. Early and intermittent suppression of SASP using JAK inhibitors alleviated chronic bone deterioration, diminished telomere dysfunction, lowered senescence and SASP marker expression, reduced bone-marrow adiposity, and mitigated radiation related lymphatic impairment. Overall, our study shows that early targeting of SASP proteins could be a potential therapeutic to prevent radiotherapy-related chronic bone loss and risk of fractures.

**Lay Summary:** Radiotherapy as part of cancer treatment is correlated with an acute increase in senescent cells. Here we show that the pro-inflammatory senescence associated secretory phenotype (SASP) becomes much more prominent soon after radiotherapy in both patients and mice. Suppression of the SASP using a Janus kinase inhibitor, ruxolitinib, reduced the overall senescent cell burden, improved bone architecture by promoting bone formation, reduced bone marrow adiposity, and mitigated radiation-induced lymphatic impairment. Overall, these findings suggest that early inhibition of the SASP may help mitigate several bone anomalies and prevent long-term bone loss and fractures after radiotherapy.

**Graphical Abstract:** 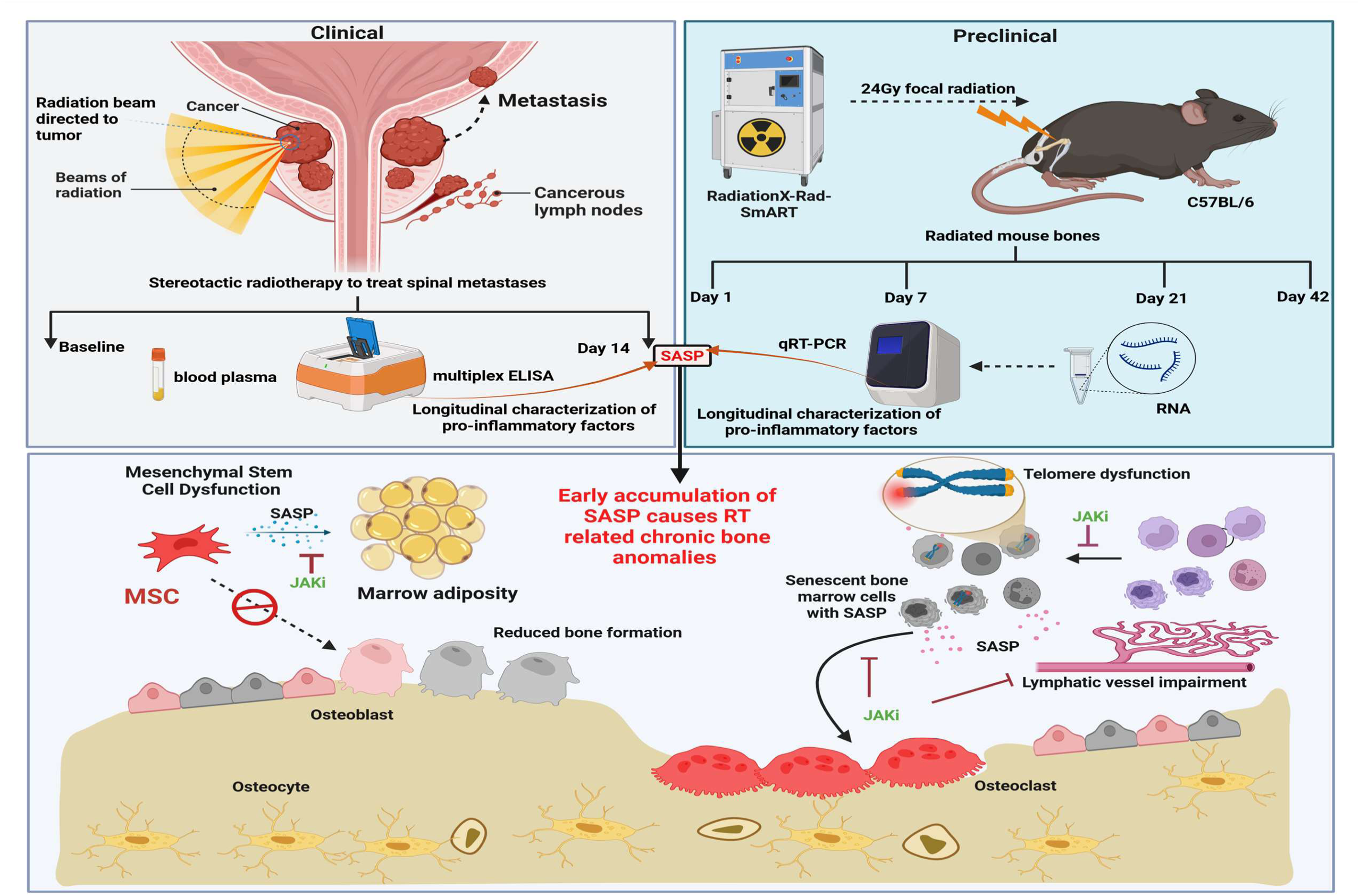

## Introduction

Growing evidence indicates that cellular senescence, not only from natural aging but also triggered by cancer treatments such as chemotherapy and radiation, is a key hallmark that promotes long term health complications and cancer related morbidities [1, 2]. Deleterious changes in the cellular secretome with aging, referred to as the senescence associated secretory phenotype (SASP), is thought to mediate the senescence burden, and to aid in the growth of cancer cells, through autocrine, paracrine and endocrine interactions. Cancer treatments such as radiotherapy and chemotherapy further exacerbate the senescent cell burden in old age.

Radiation damage to the skeleton as part of cancer treatment is a well-recognized late effect, resulting in a spectrum of bone changes from osteopenia to osteoradionecrosis [3, 4]. Unfortunately, fractures caused by radiation treatment (RT) are difficult to treat and are associated with very high rates of delayed union and nonunion [5, 6], especially in older individuals [7, 8], causing an enormous public health burden. Stereotactic body RT (SBRT) with 24Gy given as single dose [9, 10] or two daily fractions [11] is increasingly utilized to treat spinal metastases especially in oligo metastatic patients, but it is accompanied by the risk of vertebral body fractures, especially in older individuals [12–14]. Taken together, RT for cancer in the context of advancing age increases the likelihood of fractures, even with improved RT techniques. To date, the mechanism(s) underlying RT-induced bone damage has been partially elucidated [15–17], but there are no clinically proven effective treatments. Moreover, the complex mechanism underlying RT-related bone loss is still under investigation.

Clearance of senescent cells by senolytic drugs, or suppression of SASP by senomorphic drugs are an effective way to suppress age-related bone loss [18]. Prior studies from our lab have shown that senolytics alleviate RT-related bone loss [19]. It was also reported that senolytic drugs have varying abilities, and not all senolytic drugs rescued the radiated bones from architectural loss [19]. However, the pro-inflammatory SASP has not been explored as a pathophysiological mechanism underlying RT-related bone loss, nor has the therapeutic utility of agents to suppress the SASP, termed senomorphics, been tested in this clinical context.

In this study we first characterized the early pro-inflammatory protein signature in the circulation of prostate cancer patients receiving radiotherapy for spinal metastases in a longitudinal fashion. We also characterized radiated mouse bones longitudinally for the gene expression levels of pro-inflammatory SASP factors. Finally, we assessed whether early or prolonged targeting of the SASP by the senomorphic drug, ruxolitinib, a JAK/STAT pathway inhibitor (JAKi), can mitigate RT-related bone loss.

## Materials and methods

### Human plasma

All human studies were approved by the Mayo Clinic Institutional Review Board (IRB). In a single-institution prospective biobanking study, eligible patients were males with pathologically confirmed prostate cancer (either hormone-sensitive or castration-resistant) with metachronous oligometastatic disease seen on molecular imaging (≤3 extracranial metastatic lesions) and/or confirmed by biopsy (PMID: 30205124). All patients had received prior curative-intent radical prostatectomy, radiotherapy, or both prior to metastatic relapse. Patients were excluded if life expectancy was under 3 months. Plasma was collected from patients receiving RT to treat bone metastasis prior to radiation and on day 14 following radiation and assessed for 28 senescence-related proteins using ELISA.

### Senescence-related Proteins

The concentrations of 27 senescence-related proteins in plasma samples were determined with commercially available multiplex magnetic bead-based immunoassays (R&D Systems) on the Luminex xMAP multianalyte profiling platform and analyzed on MAGPIX System (Merck Millipore). The proteins quantified using this method were ADAMTS13, EOTAXIN, FAS, GDF15, GRANZYME, GROα, ICAM1, IL7, IL8, MCP1, MDC, MMP1, MMP2, MMP9, MPO, PAI1, PARC, PDGFAA, PDGFAB, RAGE, RANTES, SOST, TARC, TNFα, TNFRI, TNFRII, and VEGF (Supplementary Table S1). Activin A concentration was measured using a Quantikine ELISA Kit (R&D Systems) according to the manufacturer’s specifications. Assay performance parameters have been reported previously[20]. In cases where a variable was below the limit of detection in a sample, a value of half of the lowest measured value for that analyte was assigned.

### Animal studies

Animal studies (C57BL/6 male mice) were approved by the Institutional Animal Care and Use Committee at Mayo Clinic. All animals were housed in our facility at 23 to 25°C with a 12-h light/dark cycle and were fed with standard laboratory chow (PicoLab® Rodent Diet 20 #5053; LabDiet, St. Louis, MO, USA) with free access to water. In the past several years we have justified the use of male mice only, due to low bone mineral density in femurs of 4-month-old female C57BL/6 mice, and making it hard to assess bone loss and recovery post RT. The left femur, positioned outside the radiation field, served as a control. Following radiation, mice were administered either a vehicle (*n*=5), Dasatinib (D, 5mg/kg) + Quercetin (Q, 50mg/kg) (*n*=5) or ruxolitinib, a Janus Kinase (JAK) 1/2 specific inhibitor (JAKi; 60 mg/kg) (*n*=5) via oral gavage. Due to fighting wounds, we had to euthanize a few animals earlier than endpoint, thereby reducing the overall sample numbers as shown in the final figures. For 30Gy cumulative dosing and 60Gy cumulative dosing, groups of animals received vehicle, or D+Q once a month or JAKi thrice weekly. Animals were injected with two fluorescent labels, Alizarin (15mg/kg) and Calcein (90mg/kg), 9 and 2 days before tissue harvest at 4 months. The 24Gy single dose group received four doses of JAKi on days 1, 7, 14 and 21, and the animals were euthanized on day 42 for the various assessments (n=6/group for bone architecture, and a subset of these mice for histology and qRT PCR).

### RT-PCR

Bones were collected for mRNA isolation and qRT-PCR as described previously [19]. Briefly, a 5-mm region below the growth plate of the distal metaphyseal femur was cut out from the radiated (R) and non-radiated (NR) legs. After removal of muscle tissue, the bone samples were homogenized and total RNA was isolated using RNeasy Mini Columns (QIAGEN, Valencia, CA). cDNA was generated from mRNA using the High-Capacity cDNA Reverse Transcription Kit (Applied Biosystems by Life Technologies, Foster City, CA) according to the manufacturer’s instructions, and RT-qPCR was performed as described in our previous studies [19]. All primer sequences including senescence markers, SASP markers, osteoclast markers and adipocyte markers have been listed in Supplementary Table S2. Primers were designed so that they overlapped two exons.

### Micro-CT analysis

Bones were harvested 42-days post-focal radiation and scanned using μCT (vivaCT 40, Scanco Medical AG, Brüttisellen, Switzerland). The distal femur was scanned corresponding to a 1–5 mm area above the growth plate. All images were first smoothed by a Gaussian filter (sigma =1.2, support=2.0) and then threshold was set corresponding to 30% of the maximum available grayscale values. Volumetric bone mineral density (vBMD), bone volume fraction (BV/TV), trabecular thickness (Tb.Th), trabecular separation (Tb.Sp), trabecular number (Tb.N), and structure-model index (SMI) were calculated using 3D standard microstructural analysis.

### Histological and Immunofluorescence Methods

Immunohistochemistry was performed on tissue sections to detect lymphatic vessels (LYVE-1) and bone marrow adiposity (Perilipin). Slides were deparaffinized in xylene (3 × 5 min), rehydrated sequentially in 100%, 95%, 80%, and 70% ethanol (5 min each), and washed in deionized water (5 min). Antigen retrieval was performed using Tris-EDTA buffer (pH 9.0) at 90–95°C for 15 min, followed by cooling on ice for 15 min. Slides were washed twice with deionized water and once with PBS-TT (PBS containing 0.5% Tween-20 and 0.1% Triton X-100), followed by two washes with 1× PBS for 5 min each. Blocking was conducted for 30 min at room temperature in PBS containing 0.1% BSA and 1:60 normal goat serum. Primary antibodies against LYVE-1 (AB14917) and Perilipin-1 (D1D8) were diluted in blocking solution and incubated overnight at 4°C in a humidity chamber. Slides were washed once with PBS-TT for 5 min and twice with 1× PBS for 5 min each. Fluorescently labeled secondary antibodies specific to each primary antibody were prepared in blocking solution. Slides were incubated at room temperature for 1 h with the secondary antibody corresponding to LYVE-1 and Perilipin-1, respectively, in a sequential manner. After each secondary antibody incubation, slides were washed once with PBS-TT for 5 min and twice with 1× PBS for 5 min each. Nuclei were counterstained with DAPI, and slides were mounted using ProLong Gold Antifade mounting medium.

### TIF (Telomere Dysfunction-Induced Foci)

Deparaffinized slides were blocked with 1:60 normal goat serum in PBS containing 0.1% BSA for 30 minutes at room temperature (RT). Slides were incubated overnight at 4°C with anti-γH2A.X primary antibody (1:200, Cell Signaling, cat# 9718S) in blocking solution. Following washes in PBS-TT and PBS, slides were incubated with a biotinylated secondary antibody (1:200, Vector, cat# BA-1000) for 1 h at RT, then with Cy5 Streptavidin (1:500, Vector, cat# SA-1500) for 1 h. Slides were mounted using ProLong Gold Antifade with DAPI (Invitrogen, cat# P36935). Slides were fixed in 4% paraformaldehyde for 20 min and dehydrated in graded ethanol (70%, 90%, 100%). A hybridization mix containing Cy3-labeled PNA probe was applied (PNAbio F3002), and slides were denatured at 82°C for 10 min and hybridized at room temperature for 2 h. Excess probe was removed with formamide/SSC washes.

### Tartrate-resistant acid phosphatase (TRAP)

TRAP staining was performed using the Leukocyte Acid Phosphatase (TRAP) Kit (Sigma-Aldrich, 387A) according to the manufacturer’s protocol. Prepared TRAP staining buffer was applied to the deparaffinized slides, sealed with Parafilm, and incubated at 37°C for 2 h or overnight. Slides were counterstained with hematoxylin and imaged using a Nikon Eclipse microscope.

### Methyl methacrylate tissue embedding and dynamic histomorphometry

For the group receiving a cumulative dose of 30Gy and treated with either vehicle or JAKi, radiated and non-radiated femurs were processed at 4 months post-RT for methyl methacrylate (MMA) embedding. Dynamic histomorphometry was performed on unstained deplasticized 8-μm sections and mineral apposition rate (MAR) was calculated as described previously [21]. Dynamic histomorphometry was performed using the OsteoMeasure Histomorphometry System (OsteoMetrics, Inc., Decatur, GA, USA) with an Olympus Dotslide motorized microscope system (Olympus, Waltham, MA, USA).

### Statistical power, sample size, and computational analyses

Based on our previously published work [16, 19, 22], we estimated that a sample size of 4–6 mice per group would provide 80% power to detect a 30% reduction in BV/TV in irradiated (R) bone compared with non-irradiated (NR) contralateral bone, assuming a two-sided α of 5% (effect size 0.5; paired t test, nQuery Advisor 7.0; Statsols, Boston, MA, USA). For histological and mRNA analyses, a minimum of three mice per group has previously been sufficient to detect approximately 30% differences under a two-sided α of 5%. Sample size considerations and analytical plans were developed in consultation with the Mayo Clinic Biostatistics Core. Statistical analyses were conducted using GraphPad Prism 10.4.1 (GraphPad Software, Inc., La Jolla, CA, USA). Data are presented as medians with interquartile ranges. Unpaired two-tailed t tests were used to evaluate two groups; one-way ANOVA was used to make a comparison between three groups and two-way ANOVA followed by Dunnett’s post hoc test was applied to assess the effects of two variables and two groups. Heat maps were generated using Morpheus software (https://software.broadinstitute.org/morpheus).

## Results

### Longitudinal characterization of a pro-inflammatory secretory signature post-radiotherapy

To investigate if there are early changes in the peripheral blood in prostate cancer patients receiving radiotherapy to treat spinal metastases, blood plasma was collected at baseline and on day 14. The majority of pro-inflammatory factors assessed were elevated on day 14 after RT, including ACTIVIN A, FAS, GRO ALPHA, MCP1, IL8, MMP9, PAI1, ICAM1, MPO, TNFRI, TNFRII, PDGFAA, PDGFAB, VEGF and GDF15, as compared to day 0 (Fig. 1A, B). Conversely, TNF-ALPHA, SCLEROSTIN, MMP1 and MMP2 decreased in abundance on day 14 as compared to baseline levels (Supp. Fig.1). These data sets are also presented in a way where protein levels in individual patients could be visualized and tracked (Supp. Fig. 2 and 3).

**Figure 1.**
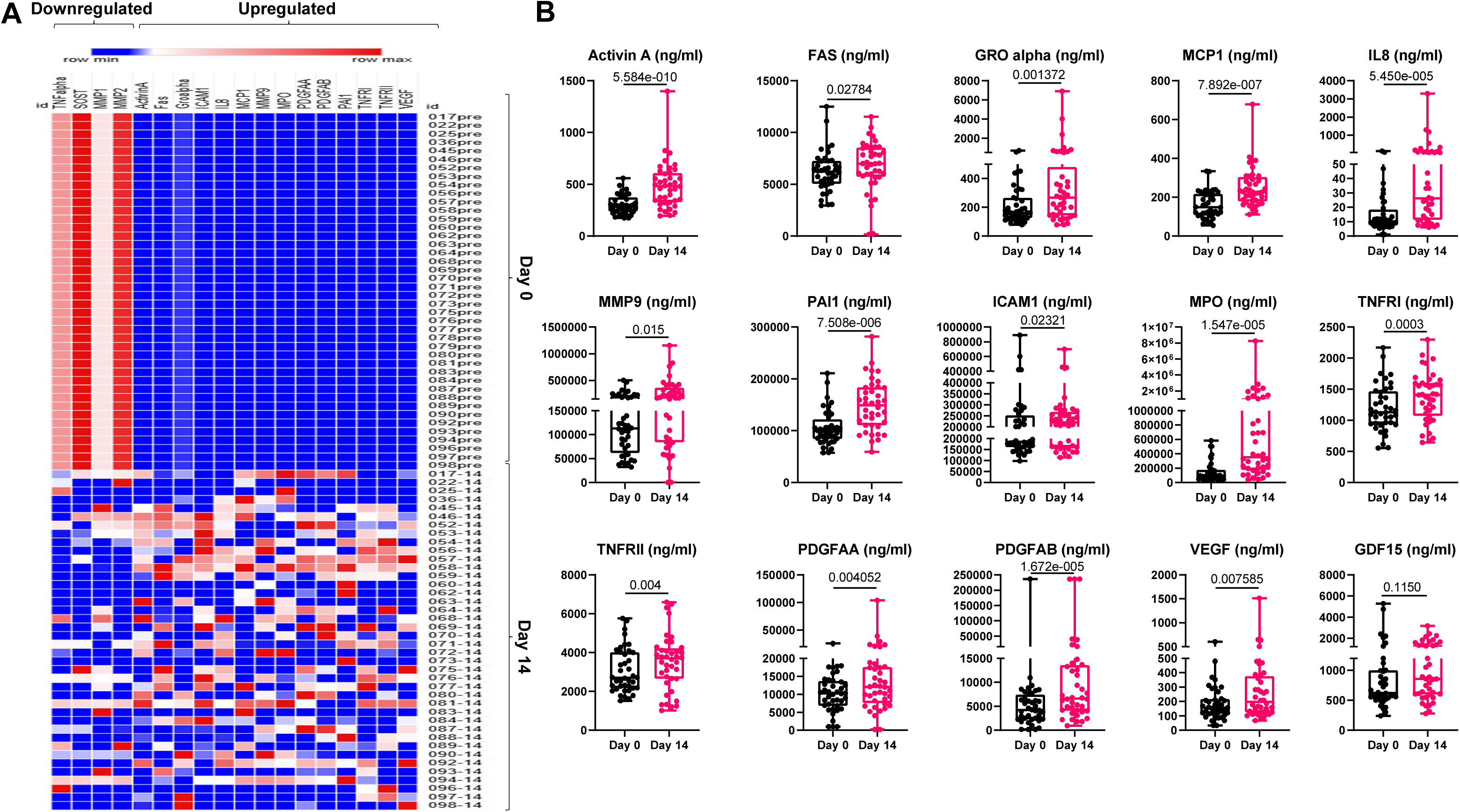
Radiation induces pro-inflammatory SASP proteins in human plasma. Human plasma samples were collected from patients receiving RT to treat spinal metastasis for evaluation of changes in pre- and post- RT protein signatures by multiplex ELISA. (A) Heatmap indicates downregulated and upregulated SASP proteins observed on day 14 as compared to day 0 in patients receiving radiotherapy. (B) Longitudinal analysis of protein levels significantly changed between pre- and post-RT. Statistical comparisons were made using a two-tailed unpaired t-test between Day 0 and Day 14.

### Longitudinal characterization of SASP genes in radiated bones

To understand if mice have similar changes in the pro-inflammatory SASP genes longitudinally as seen in our RT patient population, we performed gene expression analysis on days 1, 7, 21 and 42, in mouse femurs exposed to 24Gy focal radiation, using the non-radiated contralateral femur as the control. Majority of the SASP genes reach peak expression on day 7 in R-femurs, followed by a decline in their expression levels on days 21 and 42. *Ccl2*, *Ccl7*, *Mmp12*, and *Cxcl13* were unique in their expression profiles as they maintained their high expression levels on days 21 and 42 (Fig.2).

**Figure 2.**
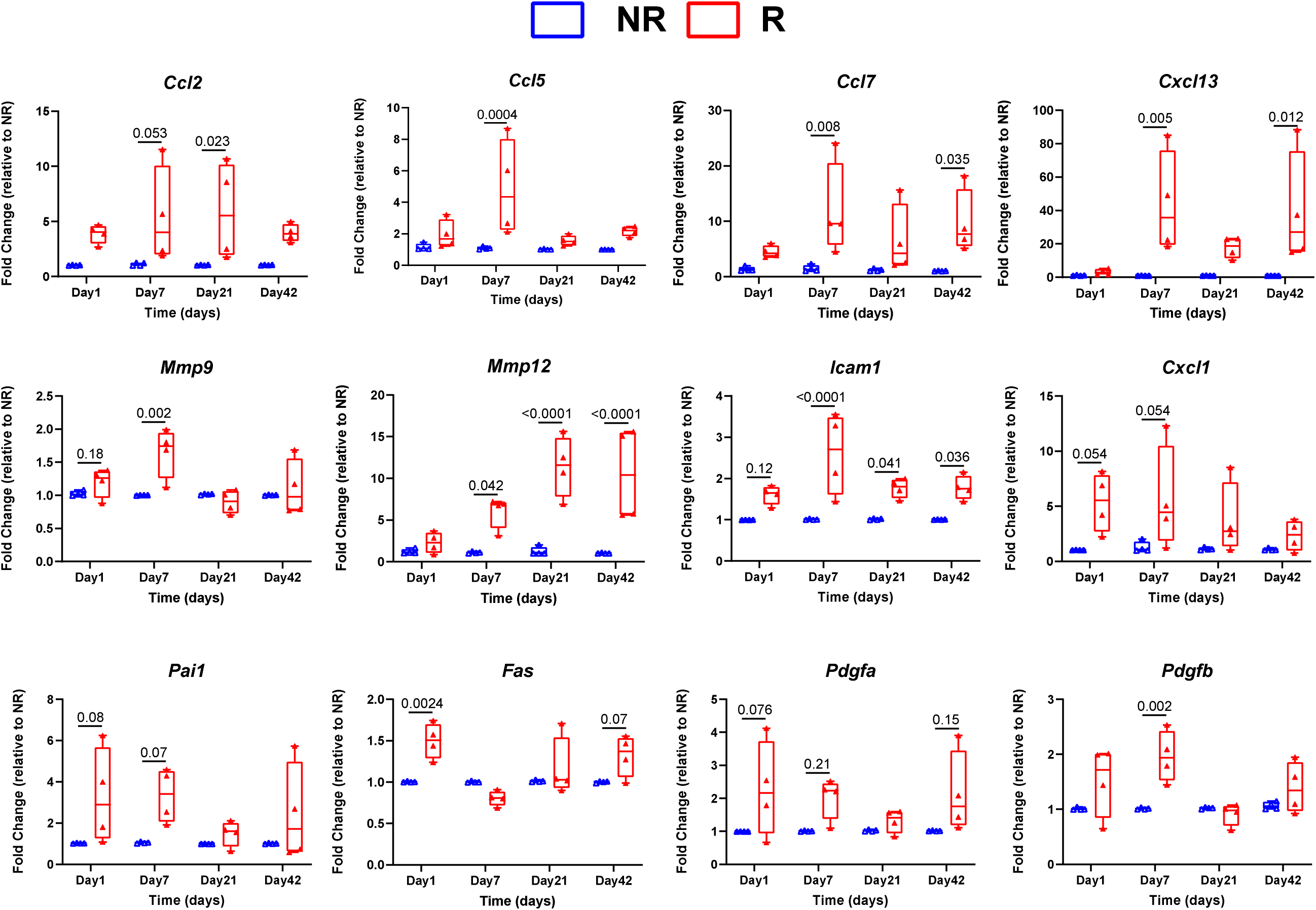
Longitudinal assessment of SASP gene expression in radiated mouse bones. The mice received a dose of 24Gy (6.6Gy /min) on day 0, delivered focally to 5mm of the femoral metaphyseal region using X-Rad-SmART. The left femur was outside the radiation area and served as control. SASP markers were measured in NR (non-radiated) and R (radiated) bones at various time points: Day 1, Day 7, Day 21, and Day 42 post-radiation. The histogram displays genes that are differentially expressed in R-bones compared to NR-bones, as detected by qRT-PCR. Fold changes are relative to NR samples. Statistical comparisons were performed using a two-way ANOVA.

### Comparative analysis between senotherapeutic agents to suppress radiation-related chronic bone damage

Previously, we showed that clearance of senescent cells by a senolytic cocktail of Dasatinib (D) and Quercetin (Q) (D+Q), administered intermittently, mitigated the 24Gy focal RT-related bone loss. Here we compared two RT dosing regimens, which model patients receiving SBRT, using cumulative dosing of RT to the 5mm area in mice femoral metaphysis, delivering either 30Gy (6Gy daily for five consecutive days) (Fig.3) or 60Gy (12Gy daily for five consecutive days) (Supp. Fig. 4). As proof of concept, we first confirmed that the senolytic cocktail, D+Q administered once a month orally, was able to significantly improve the bone architecture in the mice exposed to 30Gy after 4 months of RT, as compared to the vehicle-treated mice (Fig.3 B, C). Interestingly, with the increase in RT dose to 60Gy, D+Q failed to show the same level of efficacy in protection of bone architecture, and most of the parameters which were improved, when compared with vehicle-treated groups, became non-significant (Supp. Fig. 4).

**Figure 3.**
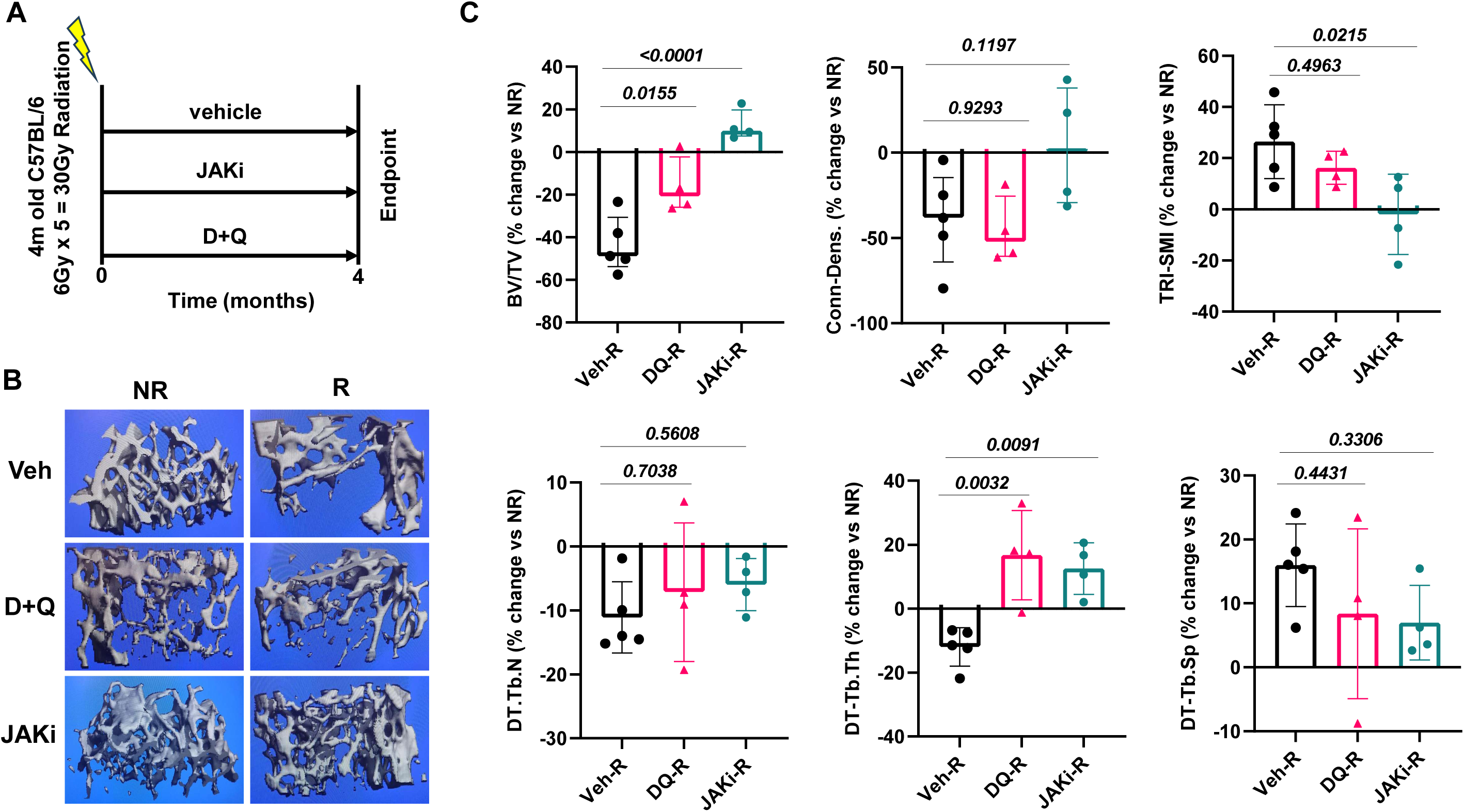
Comparative assessment of a senolytic cocktail and a senomorphic to prevent RT-related chronic bone loss. (A) A cumulative dose of 30 Gy (6 Gy × 5, administered over five consecutive days) was delivered focally to 5mm of the femoral metaphyseal region using X-Rad-SmART. Mice were treated with vehicle, JAKi, or D+Q for four months, followed by bone architecture analysis. (B) Representative three-dimensional micro-CT images indicate alterations in bone architecture among vehicle-, D+Q-, and JAKi-treated mice. (C) Micro-CT analysis to compare vehicle-, D+Q-, and JAKi-treated mice including BV/TV, Connective density, TRI-SMI, Tb. N., Tb. Th. and Tb.Sp. Statistical comparisons were performed using one-way ANOVA followed by Tukey’s multiple comparison test.

Interestingly, suppression of SASP by JAKi (administered three times a week for 4 months), was highly effective in mitigating bone loss in response to both 30Gy (Fig. 3 B, C) and 60Gy (Supp. Fig 4) RT doses. This improvement in bone architecture can be attributed largely to improvement in trabecular thickness (Tb.Th.), while no change was seen in trabecular number (Tb.N). Dynamic histomorphometry data revealed improved mineral apposition rate (MAR) in the JAKi-treated radiated bones, as compared to vehicle-treated radiated bones (Fig.4). We did not pursue the dynamic histomorphometry for the D+Q samples, as the effects of D+Q were modest and redundant to our work published previously[19].

**Figure 4.**
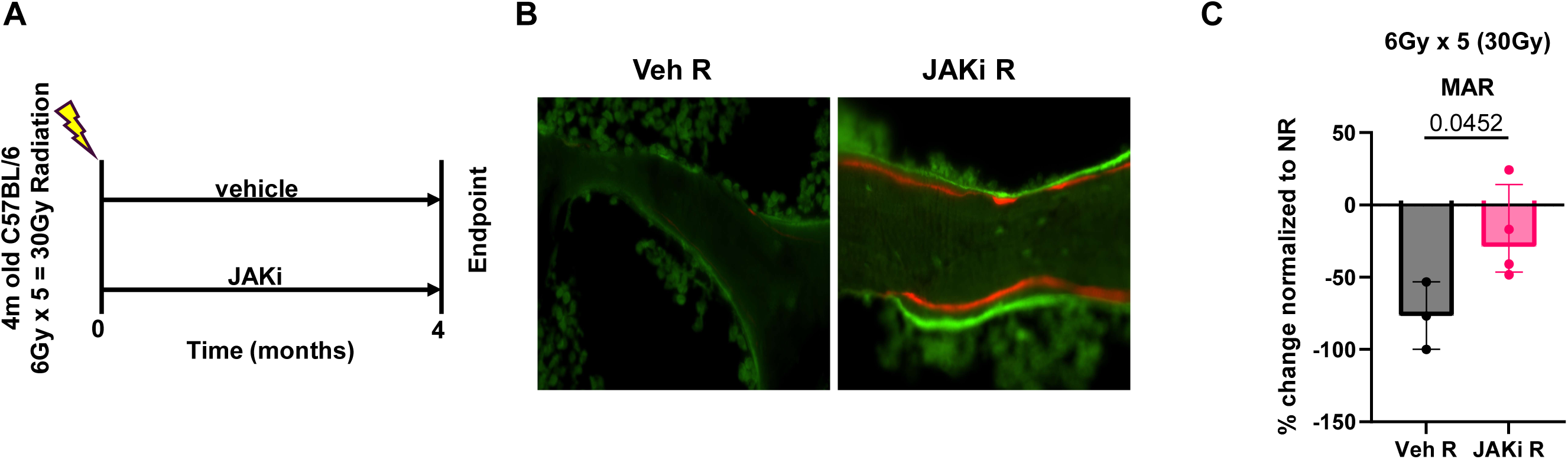
Mitigation of SASP by JAKi preserves functional osteoblasts in radiated bones. (A) A total dose of 30 Gy (6 Gy × 5) was administered over five consecutive days. Mice were treated with vehicle or JAKi for 4 months. (B) Representative images showing the double labeling of two different dyes, Alizarin and Calcein. (C) Mineral apposition rate (MAR) as a percentage change of MAR to non-radiated bones is presented. Statistical comparisons were performed using a two-tailed unpaired t-test.

### Early suppression of the SASP alleviates radiation-related chronic bone loss

Our proof-of-concept data in the 4-month animal study showed that JAKi was more effective than the senolytic approach. Since our patient and mouse data both indicated that the RT-related SASP is an early event, we further tested the efficacy of JAKi in normalizing the SASP in the short-term after RT and with a limited dose of the drug. Administering JAKi orally on days 1, 7, 14 and 21, with a gap of 21 days of no treatment before analysis, revealed a significant recovery from bone loss in mice receiving a single focal RT of 24Gy (Stereotactic Body Radiotherapy SBRT dosing regimen commonly used in clinics and in our previous studies)[19] to the 5mm area in femoral metaphysis (Fig.5). Improvements in the bone volume fraction (BV.TV) and bone mineral density (BMD) were significantly improved in the JAKi-treated radiated bones, as compared to vehicle-treated radiated bones (Fig. 5 B, C). Consistent with our 4-month chronic study, in this short-term study as well, trabecular thickening was the key change which contributed to the bone architectural improvement upon JAKi treatment.

**Figure 5.**
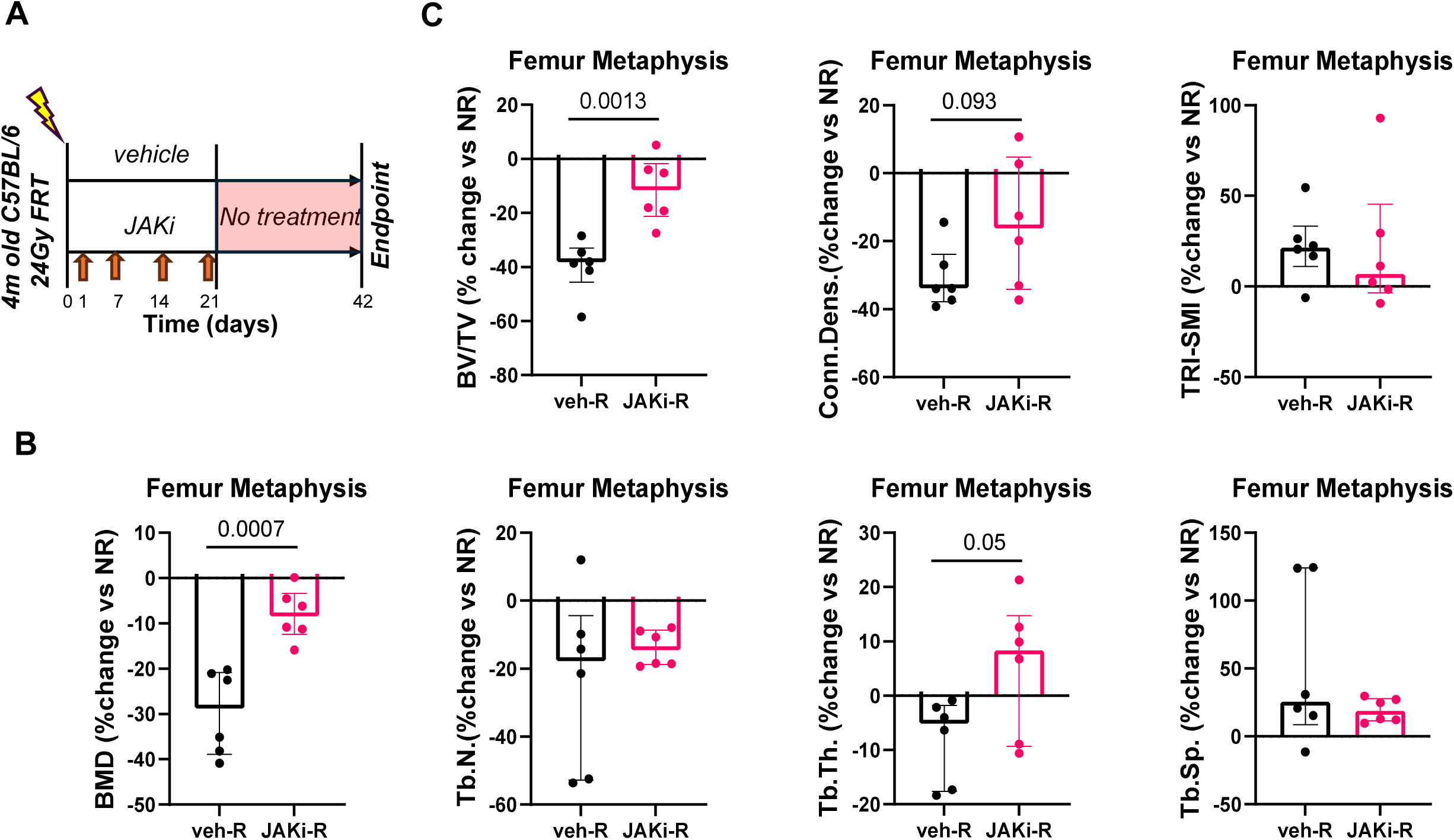
Acute suppression of SASP by JAKi modulates chronic RT-related bone architecture. The mice received a single dose of 24Gy (6.6Gy /min) on day 0, delivered focally to 5mm of the femoral metaphyseal region using X-Rad-SmART. The left femur was outside the radiation area and served as control. In a limited (early) treatment, JAKi was administered on Day 1, 7, 14, and 21, followed by 21 days of no treatment, to assess its impact on bone architecture on day 42. (B-C) Bone mineral density was analyzed using micro-CT analysis including BMD, BV/TV, Conn.Dens., TRI-SMI, Tb.N, Tb.Th. and Tb.Sp. Statistical comparisons were made using a two-tailed unpaired t-test.

### Early inhibition of the SASP suppresses radiation-related senescence and telomere dysfunction

In our model where SASP was suppressed using JAKi, but only in the early phase, with a subsequent 21-day no treatment phase, the overall senescence burden was reduced on day 42 post-RT. The cyclin dependent kinase inhibitor, p21 is a key marker of senescence if combined with telomere dysfunction and the SASP signature. Interestingly, *p21* expression was reduced significantly in the JAKi-treated radiated bones, as compared to vehicle-treated radiated bones (Fig. 6A). The expression levels are normalized to levels from the contralateral non-radiated leg. As shown previously, we saw an increase in telomere-induced dysfunction (increase in TIF, also known as telomere-associated foci, TAF) in the bone marrow cells following RT in vehicle-treated radiated bones. The TIF+ bone marrow cells were significantly lower in the JAKi-treated radiated bones (Fig. 6 B and 6C). Furthermore, to strengthen the idea that the overall senescence burden is suppressed upon JAKi treatment, we confirmed that several pro-inflammatory SASP factor genes were significantly downregulated by JAKi treatment in radiated bones as compared to vehicle-treated radiated bones.

**Figure 6.**
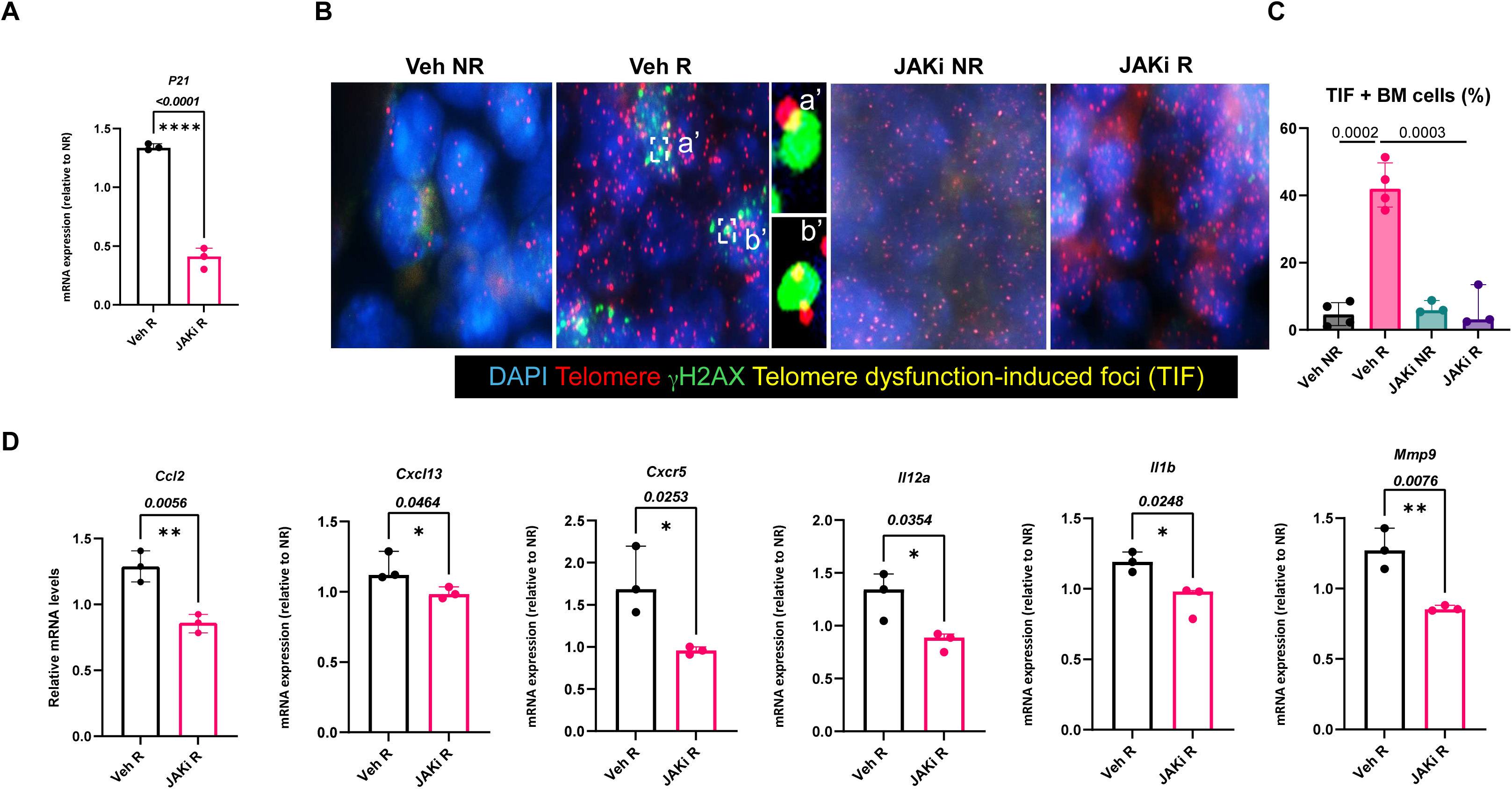
Suppression of SASP by JAKi mitigates radiation-induced senescent cell burden. The mice received a single dose of 24Gy (6.6Gy /min) on day 0, delivered focally to 5mm of the femoral metaphyseal region using X-Rad-SmART. The left femur was outside the radiation area and served as control. In a limited (early) treatment, JAKi was administered on Day 1, 7, 14, and 21, followed by 21 days of no treatment, to assess its impact on markers of senescence and SASP on day 42. (A) p21 mRNA expression levels were analyzed by RT-qPCR. Statistical comparisons were made using a two-tailed unpaired t-test. (B) Representative microscopic images indicate radiation-induced telomeric DNA damage [(shown in a’ and b’ as telomere-induced dysfunction foci (TIF), also known as telomere-associated foci (TAF)] in irradiated legs compared with vehicle-treated non-irradiated (veh-NR) control legs, between vehicle and JAKi treatment groups. (C) Quantification of TIF+ bone marrow (BM) cells, seen elevated in vehicle-treated radiated bones, and which is alleviated in the JAKi-treated radiated bones. Statistical comparisons were made using a two-way ANOVA followed by Dunnett’s post-hoc test. (D) SASP mRNA expression levels at day 42, including *Ccl2*, Cxcl13, *Cxcr5*, *Il12α*, *Il1β*, and *Mmp9*, were analyzed by RT-qPCR. Data are shown as relative expression normalized to non-irradiated controls. Statistical comparisons were made using a two-tailed unpaired t-test.

### Early suppression of SASP regulates radiation-related changes in lymphatic vessel abundance

Lymphatic vessels are reported to increase in abundance as a response to injury and stress, including radiation. As expected, the non-radiated physiological bone samples have minimal levels of lymphatic vessels in the bone marrow (Fig.7A, B). As reported by others, we were able to confirm in our vehicle-treated radiated bones that Lyve1-labeled lymphatic vessels were significantly elevated, while lymphatic vessels were dramatically reduced in the JAKi-treated radiated bones (Fig.7A, B).

**Figure 7.**
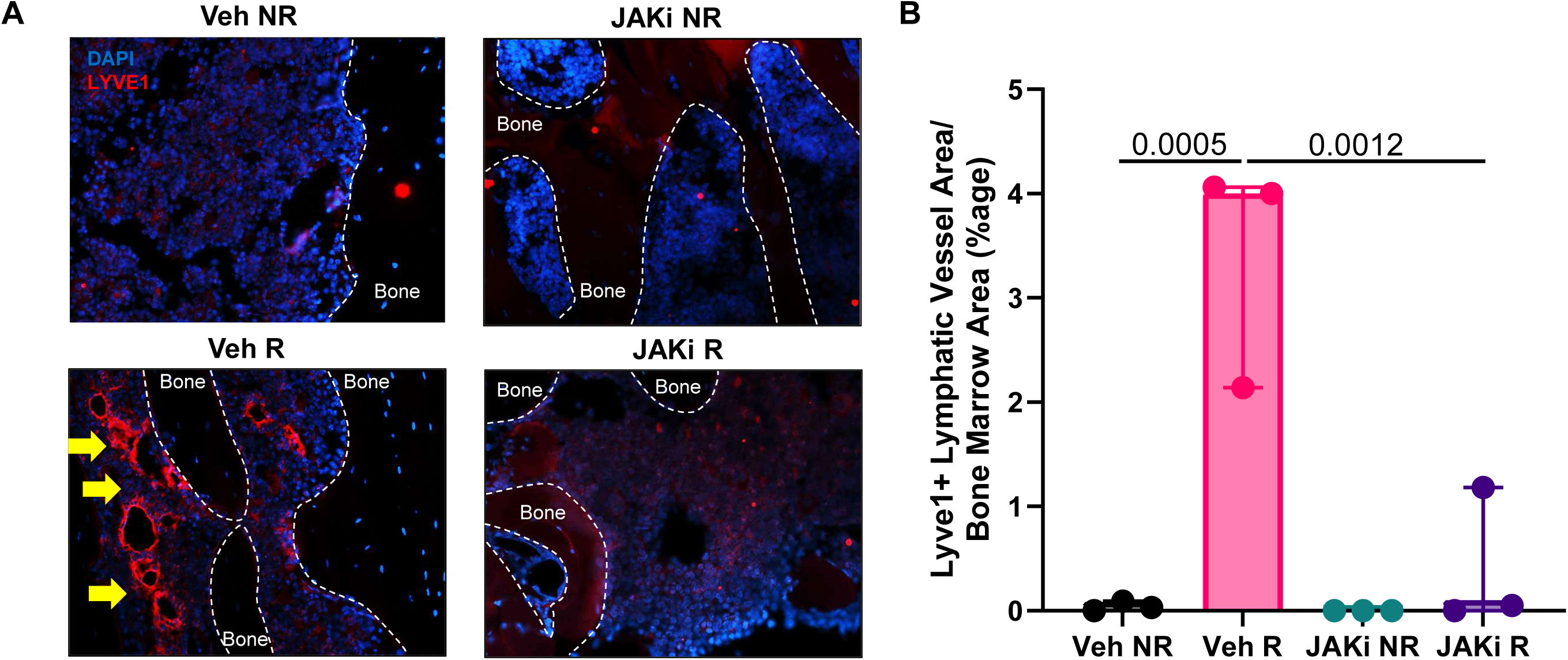
Suppression of SASP by JAKi mitigates radiation-induced bone marrow abundance of lymphatic vessels. The mice received a single dose of 24Gy (6.6Gy /min) on day 0, delivered focally to 5mm of the femoral metaphyseal region using X-Rad-SmART. The left femur was outside the radiation area and served as control. In a limited (early) treatment, JAKi was administered on Day 1, 7, 14, and 21, followed by a phase of 21 days of no treatment, to assess its impact on lymphatic vessels by immunohistochemistry on day 42. (A) Representative immunofluorescence images showing LYVE-1–positive lymphatic vessels (red) and DAPI-stained nuclei (blue) in bone marrow from the vehicle-treated or JAKi-treated groups. Bone regions are traced by the dashed lines. Yellow arrows highlight LYVE-1–positive structures. (B) Lymphatic vessel area was quantified and normalized to the bone marrow area (Bone marrow area= Total area-bone area). Statistical comparisons were made using a two-way ANOVA followed by Dunnett’s post-hoc test.

### Mitigation of SASP regulates bone marrow adiposity in radiated bones

It is well established that RT results in a lineage switching of mesenchymal stem cells (MSCs) into adipocytes. We have reported previously that a senescent bone environment triggers and maintains bone marrow adiposity (BMAd), and clearance of senescent cells by a senolytic cocktail of D+Q was able to suppress BMAd and BMAd-related genes. Here we report that suppression of SASP by JAKi could also suppress Perilipin+ BMAd and BMAd related genes such as *Cebpa*, *Igf1*, *Igf2* and *Cfd* (Fig. 8A, B).

**Figure 8.**
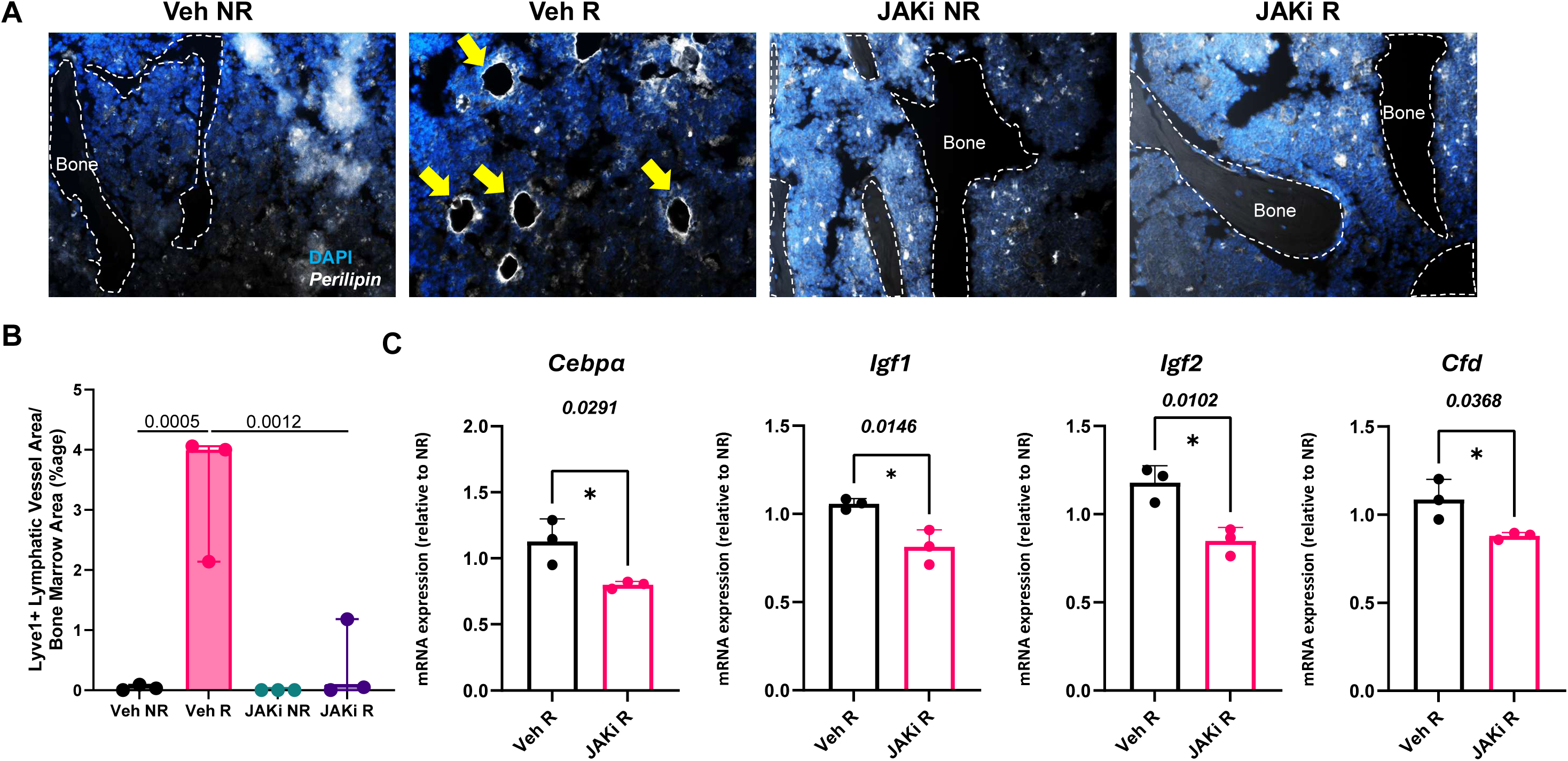
Suppression of SASP by JAKi mitigates radiation-induced bone marrow adiposity and related gene expression. The mice received a single dose of 24Gy (6.6Gy /min) on day 0, delivered focally to 5mm of the femoral metaphyseal region using X-Rad-SmART. The left femur was outside the radiation area and served as control. JAKi was administered on Days 1, 7, 14, and 21, followed by 21 days of no treatment, to assess its impact on bone marrow adiposity by immunohistochemistry on day 42. (A) Representative immunofluorescence images showing Perilipin (white) and DAPI-stained nuclei (blue) in bone marrow from vehicle-treated or JAKi-treated mice. Bone regions are traced by the dashed lines. Yellow arrows indicate Perilipin-positive marrow adipocytes. (B) Perilipin+ marrow adipocytes were quantified and normalized to the bone marrow area (Bone marrow area= Total area-bone area). Statistical comparisons were made using a two-way ANOVA followed by Dunnett’s post-hoc test. (C) Quantitative RT-qPCR analysis of *Cebpa*, *Igf1*, *Igf2*, and *Cfd* mRNA expression levels in radiated bones from vehicle-treated versus JAKi-treated groups, with the values shown as relative expression normalized to non-radiated controls. Statistical comparisons were made using a two-tailed unpaired t-test.

### Suppression of SASP inhibits osteoclasts and downregulates osteoclast-related genes

The presence of senescent cells and an associated SASP has been reported to induce increased resorption of bone by osteoclasts[18]. As compared to the vehicle-treated radiated bones, the JAKi-treated radiated bones had significantly lower TRAP+ osteoclasts (Supp. Fig.5A) and osteoclast-related genes (Supp. Fig. 5B).

## Discussion

Musculoskeletal anomalies and bone loss following RT pose significant challenge in extending the doses of RT to treat aggressive cancers [23–25]. While the exact mechanisms behind fractures post-RT are still being explored, reports have shown that a plethora of pro-inflammatory proteins are elevated [26, 27]. Here we report for the first time that the early gene expression pattern of pro-inflammatory SASP factors from radiated bones in mice, correlated with that of the pro-inflammatory proteins tested in prostate cancer patients receiving RT to treat spinal metastasis. These pro-inflammatory proteins include several cytokines, chemokines and proteases also seen in older adults [28]. As proof-of-concept to prove that the increase in SASP following RT could be the key driver for bone loss, we demonstrated in mice that the suppression of these pro-inflammatory SASP by a senomorphic drug, ruxolitinib, a JAK/STAT pathway inhibitor, was effective in improving bone architecture.

RT induces significant DNA damage in the cellular bone marrow environment including osteoblasts, osteocytes and bone marrow cells [16, 17, 29]. Some of the damaged DNA gets repaired as part of the response to stress, while a small percentage of cells undergo apoptosis. However, the majority of the cells which undergo DNA damage, largely at the telomeric sites, become senescent, activating a pro-survival pathway which allows these cells to activate the cyclin dependent kinase inhibitors (CDKIs), remain metabolic active and develop a pro-inflammatory SASP [30, 31]. Telomere shortening also leads to a senescent state [32, 33], however telomeric DNA damage (TIF/ TAF) and not telomere shortening is the main cause of radiation-induced senescence of cells [34]. Reversing telomere shortening by telomerase catalytic subunit was able to prevent replicative senescence in human osteoblasts [35], while telomerase activity restoration could restore human skin from radiation-induced DNA damage [36]. Over time, senescent cells accumulate gradually, with clearing by the immune system until the immune response becomes blunted by aging and these senescent cells increase in abundance with a robust SASP. RT not only generates DNA damage several fold higher as compared to aging, but does that during a short span of time, generating a high senescent cell burden with local and systemic effects [22]. This high senescence environment was clearly reflected by the presence of several SASP proteins on day 14, in the circulation of patients following RT. Whether SASP directly leads to bone loss remains to be explored, studies have demonstrated that SASP can elevate osteoclastogenesis and increase the level of bone resorption [18].

Suppression of the SASP by senomorphic drugs has been effective in suppressing age-related bone loss [18], skin aging [37], in age-related changes in senescent retinal pigmental cells [38] and in mitigation of cancer cachexia[39]. Moreover, the nature of these senomorphic drugs depend on the pathways they target but overall seem to be an effective way to suppress the SASP without affecting the senescent cells. However, suppression of the SASP can also prevent the generation of new senescent cells by inhibiting autocrine, paracrine or endocrine signaling of inflammatory and other factors. This was confirmed in our data, where suppression of SASP by JAKi reduced levels of *p21* expression and TIF+ senescent bone marrow cells, providing the first evidence that suppression of SASP can ameliorate senescent cell burden in the bone microenvironment after irradiation.

The comparatively improved efficacy with the senomorphic, JAKi, as compared to the senolytic approach of D+Q, was surprising, but potentially can be explained by the way these drugs work. Senescent cells are heterogenous in nature based on triggers that made them senescent and the mechanisms by which they become resistant to apoptosis. Since there is specificity between an individual senolytic drug and the anti-apoptotic pathway that it targets, not all senolytic agents will target all senescent cells [40–43]. Conversely, while NF-КB, p38 MAPK, cGAS-STING and mTOR also play critical roles in regulating different facets of the SASP, the JAK/STAT signaling pathway is considered the hub for a majority of inflammatory SASP proteins [44–48]. While D+Q and JAKi both were able to show an improved bone architecture in the cumulative 30Gy radiated femurs, as compared to the non-radiated controls, JAKi was indeed more effective in cumulative 60Gy dosing than D+Q, which failed to rescue the bone loss after this regimen. Further investigation of the JAKi efficacy at 30Gy revealed an improved mineral apposition rate as compared to the vehicle treated mice. As shown previously, several regions of the vehicle-treated radiated femurs showed little to no labeling, indicating the complete suppression of bone formation and supporting the plausible mechanism by which JAKi preserves bone architecture.

Another aspect we focused on in this study was understanding the early changes post-RT that may be responsible for the chronic bone damage and fractures. It has been reported that post-RT, hematologic changes are observed in the early phase often correlating with the chronic toxicity of organs [49]. Our clinical and preclinical data shown here strongly suggests that there is an elevated pro-inflammatory signature present within the first few days and weeks post-RT, potentially responsible for chronic bone loss. To test this, limited and early suppression of SASP by JAKi was able to prevent chronic bone loss in animal models of RT. Further studies are required to demonstrate if this treatment window can be further reduced to the first week post-RT only (in both patients and pre-clinical models), but our data is proof of concept that an early intervention targeting the pro-inflammatory SASP could be a potential treatment strategy to prevent potential bone loss and fractures due to RT.

RT also causes stress which results in yellowing of the marrow, which at cellular level is due to lineage switching of the MSCs into adipocytes, causing accumulation of BMAd [16]. Suppression of SASP by JAKi showed lower BMAd levels, which is confirmation of the hypothesis that a high senescence burden is linked to high BMAd levels [22]. Following radiation injury (genotoxic stress), lymphatic vessels within the bone undergo significant expansion and proliferation, potentially assisting in the regeneration process [50]. Following exposure to radiation we saw a similar increase in Lyve1+ lymphatic vessels in bone, while suppression of SASP by JAKi reduced the accumulation of lymphatic vessels. While our data cannot confirm conclusively, this reduction in lymphatic vessels may indicate that the regeneration process was completed early or that it was obviated by the early inhibition of the SASP. In the absence of any treatment, RT-related injury does not get resolved in a clinically meaningful manner and delays bone regeneration.

Overall, our study provides the first indication that elevated circulatory pro-inflammatory proteins could be derived from bone tissue acutely exposed to RT. Furthermore, targeting this early SASP signature using a senomorphic drug presents a potentially viable therapeutic option to prevent chronic bone loss by suppressing the overall senescence burden mediated by the SASP, reducing the accumulation of marrow fat and promoting early resolution of bone injury.

## Author Contributions

**David Achudhan:** Experimentation and Data collection and analysis; writing – review and editing. **Jacob Orme:** Data collection and analysis; funding acquisition; writing – review and editing. **Ritika Sharma**: Data collection and analysis, writing – review and editing. **Aqsa Komel**: Data collection and analysis, writing – review and editing. **Khubaib Gul Khan**: Data collection and analysis, writing – review and editing. **Nathan K. LeBrasseur**: Data collection and analysis, writing – review and editing. **Sundeep Khosla:** funding acquisition, writing – review and editing. **Sean S. Park:** funding acquisition; writing – review and editing. **Robert Pignolo:** Conceptualization; project administration; resources; writing – original draft; writing – review and editing; **Abhishek Chandra:** Conceptualization; project administration; funding acquisition; resources; data analysis; writing – original draft; writing – review and editing.

## Acknowledgements

This work was made possible by the Robert and Arlene Kogod Professorship (to RJP), R01 AG082681 (A.C), P01 AG062413 (S.K.), R01 AG086085 (S.K) R01 AG076515 (S.K.), Hevolution HR-GRO-23-1199144-8 (S.K), K12 CA90628-23 (J.J.O.), AG R01055529 (N.K.L.) and Eagle’s Cancer Telethon Fund for Cancer (A.C., S.P., J.J.O.)

---

**Supplementary Figure 1.**
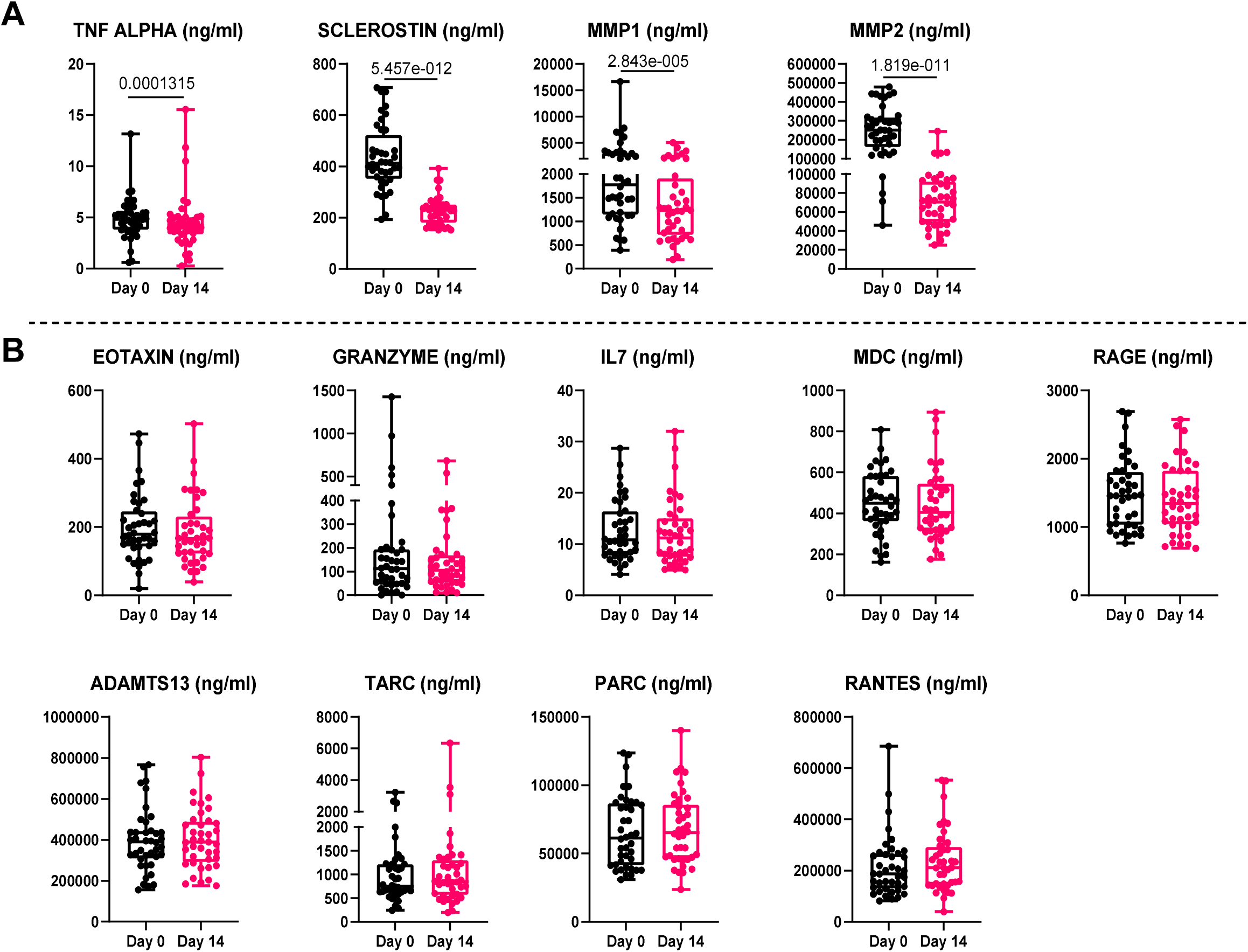

**Supplementary Fig.2.**
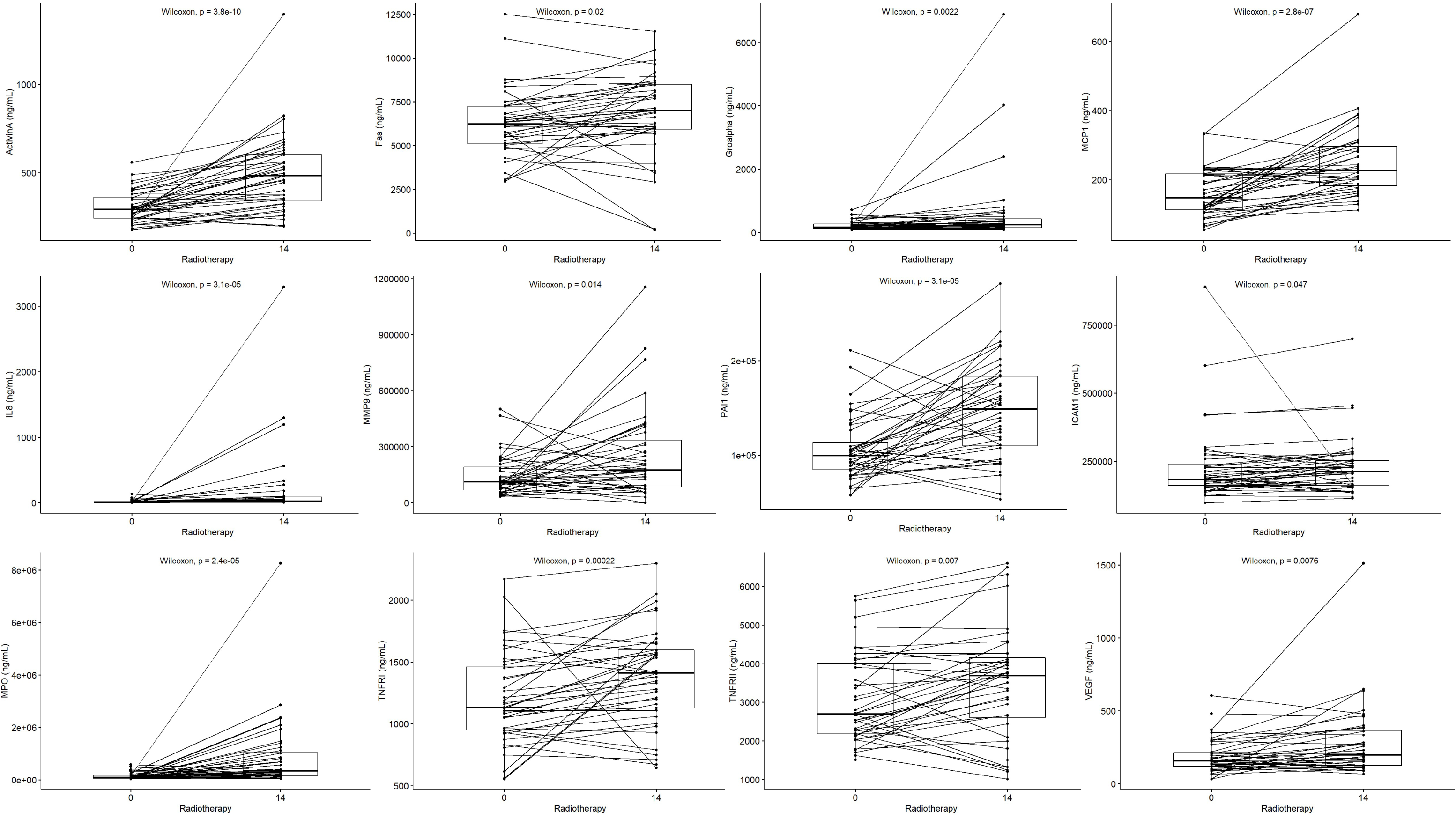

**Supplementary Fig. 3.**
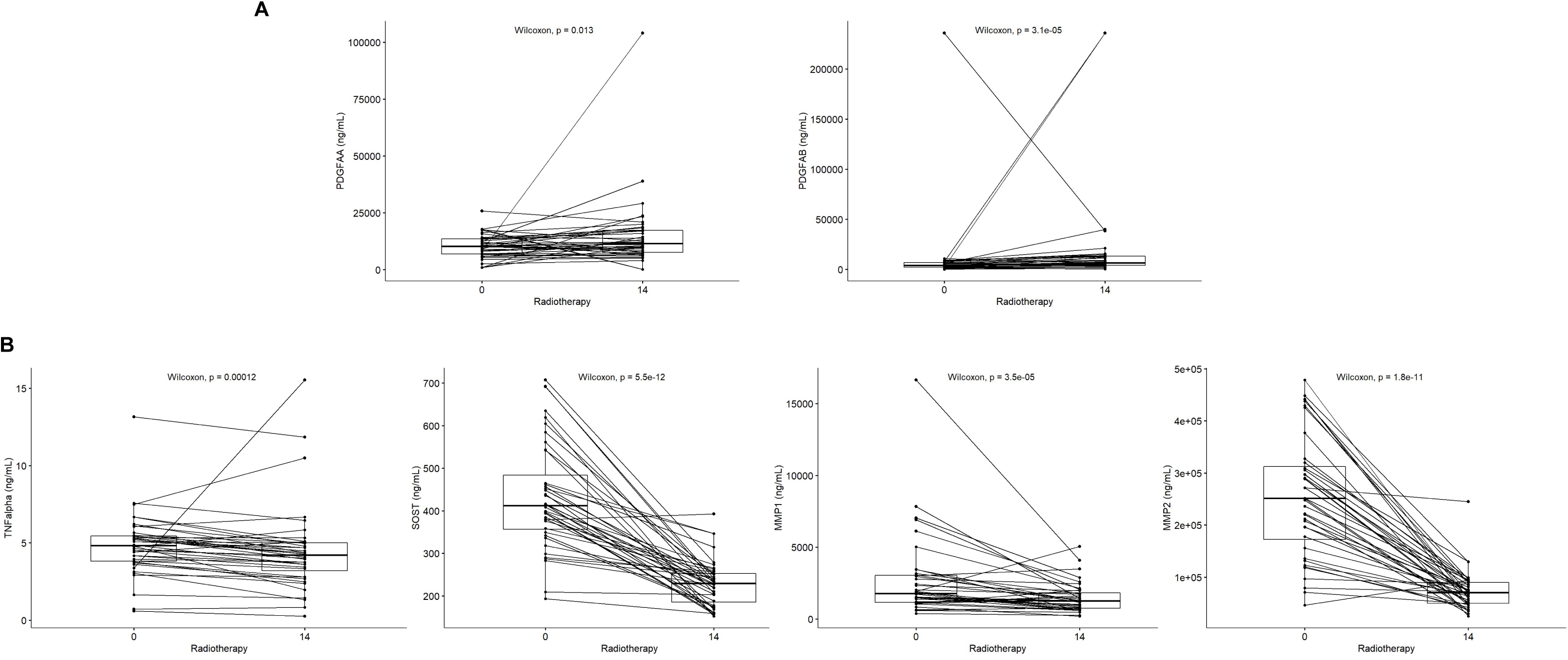

**Supplementary Fig. 4.**
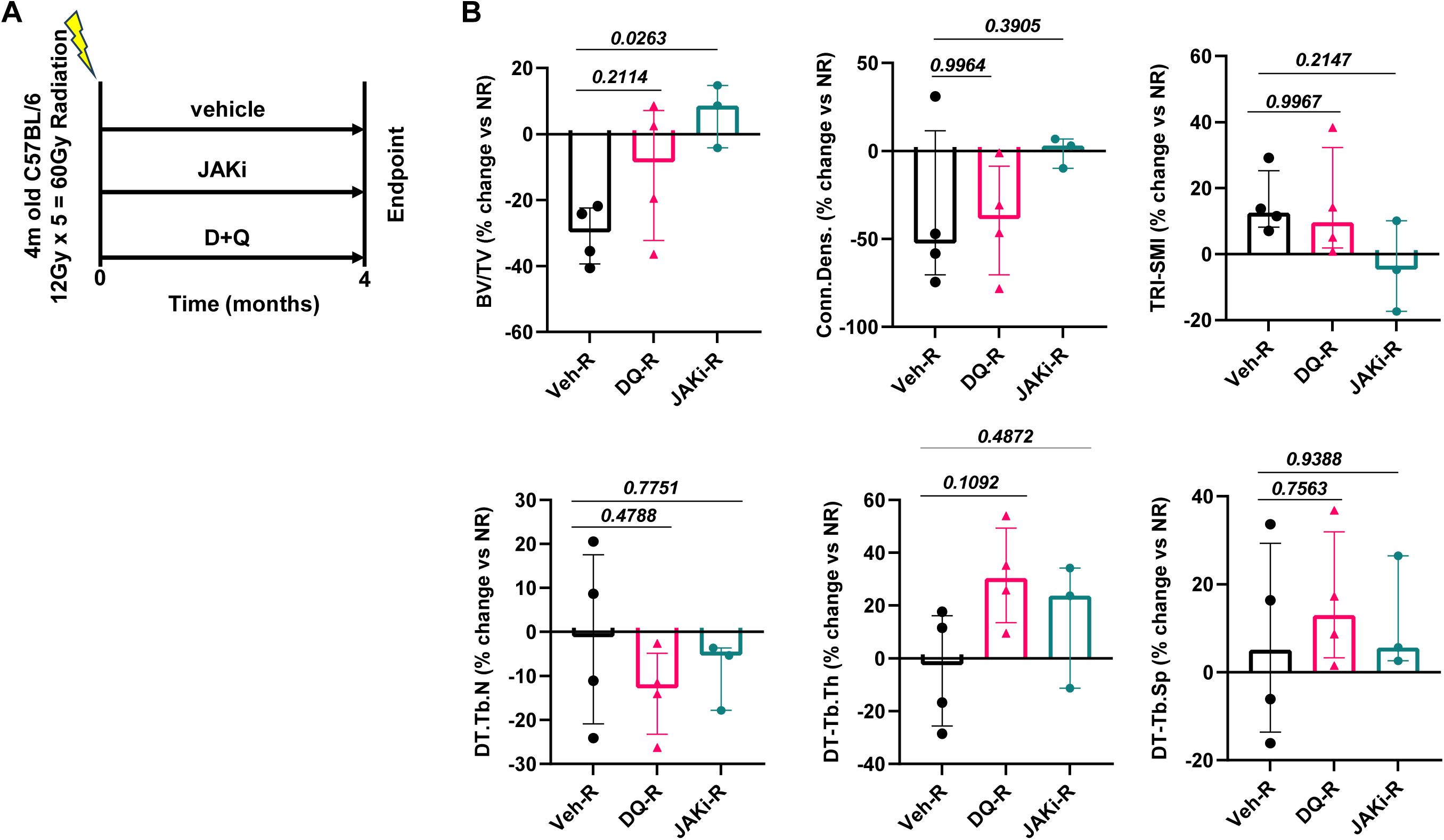

**Supplementary Figure 5.**
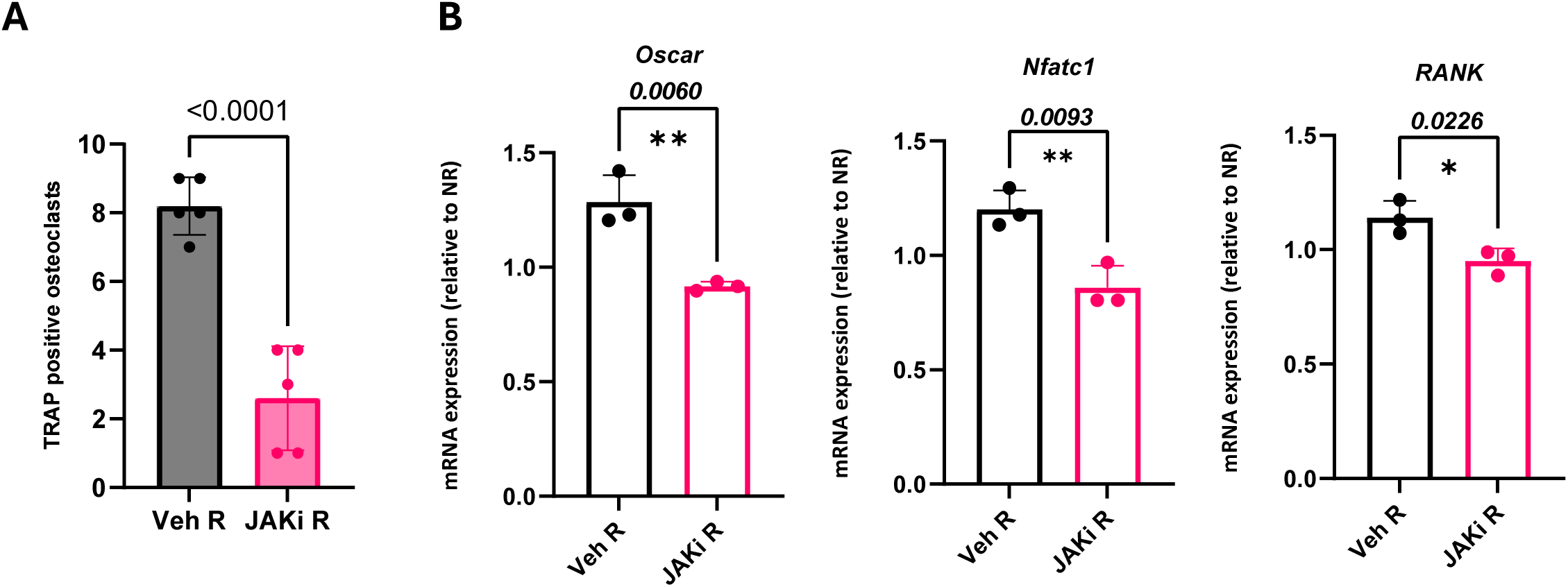

