## Supplementary material for "Suppression of early pro-inflammatory senescent signature post-radiotherapy mitigates chronic bone damage": Supp. Fig. and table

**Supplementary Figure Legend**

**Supplementary Figure 1. Changes in pro-inflammatory SASP proteins in human plasma following radiotherapy.** Human plasma samples were collected from patients receiving RT to treat spinal metastasis for evaluation of changes in pre- and post- RT protein signatures by multiplex ELISA. Longitudinal analysis of protein levels significantly downregulated (A) and proteins tested but those that did not change (B) post-RT. Statistical comparisons were made using a two-tailed unpaired t-test between Day 0 and Day 14.

**Supplementary Figure 2. Longitudinal tracking of upregulated pro-inflammatory SASP proteins in human plasma following radiotherapy.** Human plasma samples were collected from patients receiving RT to treat spinal metastasis for evaluation of changes in pre- and post- RT protein signatures by multiplex ELISA. Longitudinal analysis of protein levels significantly upregulated post-RT is shown comparing the same patient values on Day 0 and Day 14. Statistical comparison was made using a Wilcoxon signed-rank test.

**Supplementary Figure 3. Longitudinal tracking of additional upregulated and downregulated pro-inflammatory SASP proteins in human plasma following radiotherapy.** Human plasma samples were collected from patients receiving RT to treat spinal metastasis for evaluation of changes in pre- and post- RT protein signatures by multiplex ELISA. Longitudinal analysis of protein levels significantly upregulated (A) and significantly downregulated (B) post-RT. Statistical comparison was made using a Wilcoxon signed-rank test comparing the same patient values on Day 0 and Day 14.

**Supplementary Figure 4. Comparative assessment of a senolytic cocktail and a senomorphic to prevent 60Gy RT-related chronic bone loss**. (A) A cumulative dose of 60 Gy (12Gy × 5, administered over five consecutive days) was delivered focally to 5mm of the femoral metaphyseal region using X-Rad-SmART. Mice were treated with vehicle, JAKi, or D+Q for four months, followed by bone architecture analysis. (B) Micro-CT analysis to compare vehicle-, D+Q-, and JAKi-treated mice including BV/TV, Connective density, TRI-SMI, Tb. N., Tb. Th. and Tb.Sp. Statistical comparisons were performed using one-way ANOVA followed by Tukey’s multiple comparison test.

**Supplementary Figure 5. Suppression of SASP by JAK1/2 inhibitor mitigates radiation-induced osteoclastogenesis and associated markers.** The mice received a single dose of 24Gy (6.6Gy /min) on day 0, delivered focally to 5mm of the femoral metaphyseal region using X-Rad-SmART. The left femur was outside the radiation area and served as control. JAKi was administered on Day 1, 7, 14, and 21, followed by 21 days of no treatment, to assess its impact on TRAP+ osteoclasts by immunohistochemistry and on osteoclast-related genes on day 42. (A) Quantification of TRAP-positive cells in vehicle-treated irradiated (Veh R) and JAKi-treated irradiated (JAKi R) groups. (B) Quantitative RT-qPCR analysis of *Oscar*, *Nfatc1*, and *Rank* mRNA expression levels in irradiated vehicle-treated (Veh R) and irradiated JAKi-treated (JAKi R) groups. Data are shown as relative expression normalized to non-irradiated controls. Statistical comparisons were made using a two-tailed unpaired t-test.

**Supplementary Table S1:** List of the Proteins Measured in the Plasma of CALERIE^TM^ Participants

| **Protein name** | **Protein full name** | **Alias** |
| --- | --- | --- |
| Activin A | Activin A | INHBA |
| ADAMTS13 | A disintegrin and metalloproteinase with thrombospondin motifs 13 | VWFCP |
| Eotaxin | Eotaxin | CCL11 |
| Fas | Tumor necrosis factor receptor superfamily member 6 | APT1, TNFRSF6 |
| GDF15 | Growth/differentiation factor 15 | MIC1, NAG1, NRG1 |
| ICAM1 | Intercellular adhesion molecule 1 | CD54 |
| IL7 | Interleukin 7 |  |
| IL8 | Interleukin 8 | CXCL8 |
| MCP1 | Monocyte Chemoattractant Protein-1 | CCL2 |
| MDC | Macrophage-derived chemokine | CCL22, SCYA22 |
| MMP1 | Matrix metalloproteinase 1 | Interstitial collagenase |
| MMP2 | Matrix metalloproteinase 2 | CLG4A |
| MMP9 | Matrix metalloproteinase 9 | CLG4B |
| MPO | Myeloperoxidase |  |
| PAI1 | Plasminogen activator inhibitor 1 | SERPINE1, PLANH1 |
| PARC | Pulmonary and activation-regulated chemokine | CCL18 |
| PDGFAA | Platelet-Derived Growth Factor-AA |  |
| PDGFAB | Platelet-Derived Growth Factor-AB |  |
| RAGE | Advanced glycosylation end product-specific receptor |  |
| RANTES | Regulated on Activation, Normal T Cell Expressed and Secreted | CCL5, SCYA5 |
| SOST | Sclerostin | DAND6 |
| TARC | Thymus and activation-regulated chemokine | CCL17, SCYA17 |
| TNFα | Tumor necrosis factor alpha | TNFSF2 |
| TNFR1 | Tumor necrosis factor receptor 1 | TNFRSF1A, CD120a |
| TNFR2 | Tumor necrosis factor receptor 2 | TNFRSF1B |
| VEGF | Vascular endothelial growth factor | VPF |

**Supplementary Table S2:** List of primers were used in this study

| **Marker** | **Gene** | **Forward Primer sequence** | **Reverse Primer sequence** |
| --- | --- | --- | --- |
| SEN | *Cdkn1a* | GAACATCTCAGGGCCGAAAA | TGCGCTTGGAGTGATAGAAATC |
| SASP | *Ccl2* | AGGTGTCCCAAAGAAGCTGT | ACAGAAGTGCTTGAGGTGGT |
|  | *Ccl5* | GCCCACGTCAAGGAGTATTTCT | ACAAACACGACTGCAAGATTGG |
|  | *Ccl7* | CCCTGGGAAGCTGTTATCTTCA | CTGATGGGCTTCAGCACAGA |
|  | *Cxcl1* | CCGAAGTCATAGCCACACTCAA | CAAGGGAGCTTCAGGGTCAAG |
|  | *Cxcl13* | AGCACAGCAACGCTGCTTCT | AATACCGTGGCCTGGAGAGA |
|  | *Cxcr5* | CCTTCGCTGGCGTAAAGTTC | ACAGCCCAGCTTGGTCAGAA |
|  | *Fas* | ACCTCCAGTCGTGAAACCAT | CTCAGCTGTGTCTTGGATGC |
|  | *Icam1* | TGCTTTGAGAACTGTGGCAC | GCTCAGTATCTCCTCCCCAC |
|  | *Il12a* | ATCCTGCTTCACGCCTTCAG | GATAGCCCATCACCCTGTTGA |
|  | *Il1b* | TCAGGCAGGCAGTATCACTCA | CACGGGAAAGACACAGGTAGCT |
|  | *Fas* | ACCTCCAGTCGTGAAACCAT | CTCAGCTGTGTCTTGGATGC |
|  | *Mmp9* | AAAACCTCCAACCTCACGGA | GTGGTGTTCGAATGGCCTTT |
|  | *Mmp12* | GTGCCCGATGTACAGCATCTT | GGTACCGCTTCATCCATCTTG |
|  | *Pai1* | CAATGACTGGGTGGAAAGGC | GGGGTGGTGAACTCAGTGTA |
|  | *Pdgfa* | ATCGGCCAACTTCCTGATCT | ATCTGTCTCCTCCTCCCGAT |
|  | *Pdgfb* | CCGGTCCAGGTGAGAAAGAT | AACTTTCGGTGCTTGCCTTT |
| Osteoclast | *Nfatc1* | GAGACAGACATCGGGAGGAAGA | GTGGGATGTGAACTCGGAAGA |
|  | *Oscar* | CTCTTCAAAAGTGGCCTTGTCA | GGAAGAACTCAGCCAGCTCAA |
|  | *Tnfrsf11a* | ACTGAGGAGACCACCCAAGGA | TGAAGAGGAGCAGAACGATGAG |
| Adipocyte | *Cebpa* | GAGCCGAGATAAAGCCAAACA | CGGTCATTGTCACTGGTCAACT |
|  | *Cfd* | TGTCAATCATGAACCGGACAA | GGTGACTACCCCGTCATGGT |
|  | *Igf1* | AAAAGCAGCCCGCTCTATCC | CTTCTGAGTCTTGGGCATGTCA |
|  | *Igf2* | CATCGTCCCCTGATCGTGTTA | TTGCTGGACATCTCCGAAGAG |
| *Controls* | *βactin* | AATCGTGCGTGACATCAAAGAG | GCCATCTCCTGCTCGAAGTC |
|  | *Tuba1* | GGTTCCCAAAGATGTCAATGCT | CAAACTGGATGGTACGCTTGGT |
